# The ABCs of Cyanobacterial Glycogen: *In Vitro* Modelling of Glycogen Synthesis and Functional Divergence of Glycogen Synthases in *Synechocystis* sp. PCC 6803

**DOI:** 10.1101/2025.11.13.688228

**Authors:** Kenric Lee, Dimitrios Bekiari, Sofia Doello, Karl Forchhammer

## Abstract

Glycogen is the principal carbon reserve in *Synechocystis* sp. PCC 6803. We reconstituted its biosynthetic pathway in vitro—GlgC (Glucose-1-phosphate adenylyltransferase), two glycogen synthase isoenzymes (GlgA1, GlgA2) and the branching enzyme GlgB—to define how supply, polymerisation and branching set flux and product structure. GlgA2 shows higher specific activity and cooperates with GlgB-generated branched primers, whereas GlgA1 has higher substrate affinity and responds more to primer concentration. Product profiling links mechanism to architecture: GlgA1 produces more-branched glycogen, while GlgA2 yields longer, less-branched polymers, with GlgB biasing utilisation towards GlgA2. The complementary behaviours of GlgA1 and GlgA2 provide capacity for rapid accumulation versus steady-state maintenance and offer dynamic metabolic levers to tune glycogen content and architecture in cyanobacteria.

## Introduction

Cyanobacteria are a large group of photoautotrophic prokaryotes also capable of heterotrophic growth. Owing to their adaptability, representatives of cyanobacteria have been found to be successful in nearly every biome on the planet and are regarded as the precursors to higher plants [1]. Glycogen is the principal carbon storage polysaccharide in cyanobacteria, playing a central role in cellular energy homeostasis and adaptation to fluctuating environmental conditions [2–4].

In *Synechocystis* sp. PCC 6803 (hereafter *Synechocystis*), the pathway for glycogen biosynthesis is orchestrated by a suite of specialized enzymes [5]. Glycogen synthesis begins with the starting molecule, glucose-1-phosphate (Glc-1P) which is interchangeably converted from glucose-6-phosphate (Glc-6P) mainly by phosphoglucomutase (PGM, product of *sll0726*, EC:5.4.2.2). Glc-1P is then utilised in the initial, rate-limiting step catalysed by Glucose-1-phosphate adenylyltransferase (GlgC, product of *slr1176*, EC:2.7.7.27), which synthesizes ADP-glucose (ADP-Glc) from ATP and Glc-1P—a process subject to intricate allosteric regulation by metabolic effectors such as 3-phosphoglycerate (3-PGA) and inorganic phosphate (Pi) [6–9]. The ADP-Glc generated then serves as the activated glucosyl donor for glycogen synthase (GlgA) enzymes (EC:2.4.1.21), which directly catalyse the elongation of a growing glycogen chain. These processes mirror the complexity observed in higher plants and shed light on evolutionary parallels in α-glucan metabolism [5]

*Synechocystis* encodes two distinct GlgA isoenzymes, GlgA1/*sll0945* and GlgA2/*sll1393*. Unlike most bacteria, which typically possess a single GlgA, these isoforms display complementary and, in some contexts, specialized roles in glycogen polymerization and chain elongation [5]. Recent studies indicate that GlgA1 and GlgA2 differ in their chain extension properties—GlgA1 being progressive and GlgA2 more distributive—affecting the fine structure and branching patterns of the resulting glycogen granules [5,10]. These differences suggest isoenzyme divergence for nuanced physiological or metabolic adaptation and underscore the need for a systematic characterization of their respective biochemical properties. The final step in the maturation of glycogen involves the glycogen branching enzyme (GlgB/*sll0158* (EC:2.4.1.18), which inserts α-1,6-glycosidic linkages, increasing the number of non-reducing ends and enhancing the water solubility of the polymer [11,12]. The interplay between GlgA and GlgB is a key determinant of the overall structure, size, and accessibility of glycogen particles, which in turn can influence the cell’s capacity for rapid carbon mobilization under stress or during recovery from nutrient deprivation [13,14].

Despite these advances, understanding of how GlgAisoenzymes coordinate with GlgC and GlgB remains incomplete. Questions persist regarding their kinetic properties, primer utilization preferences, differential regulation by allosteric metabolites, and the structural outcomes of their activity under varying enzymatic and substrate conditions. Dissecting these parameters is critical for comprehending not only glycogen metabolism in cyanobacteria but also its parallels with starch biosynthesis in plants and the broader evolutionary context of polysaccharide storage in photoautotrophic organisms.

In this work, we establish an integrated *in vitro* model for *Synechocystis* glycogen synthesis, leveraging precise biochemical assays to dissect the individual and collective roles of GlgA1, GlgA2, and GlgB. Our work aims to clarify substrate selectivity, primer dependency, regulatory mechanisms, and product structure, providing new insights into the functional divergence of GlgA isoenzymes and their physiological significance.

## Results

### Comparable catalytic efficiencies of GlgA1 and GlgA2

We first revisited the kinetic properties of the GlgA isoenzymes by initiating glycogen synthesis with direct addition of ADP-Glc and using bovine glycogen as the primer (Assay A) GlgA activity was monitored via a well-documented coupled spectrophotometric assay detecting ADP release through pyruvate kinase (PK) and lactate dehydrogenase (LDH) activities in the presence of phosphoenolpyruvate (PEP) and NADH (**Figure 1**) [6,15]. As shown in **Figure 2A**, GlgA2 (0.77 ± 0.02 U mg⁻¹) exhibited approximately 50% higher activity than GlgA1 (0.51 ± 0.01 U mg⁻¹), albeit with a slightly higher K_m_ for ADP-Glc (GlgA2 K_m_: 219 ± 9.9 µM; GlgA1 K_m_: 152 ± 3.6 µM). The resulting catalytic efficiency (K_cat_/K_m_) of GlgA1 was calculated to be 3.2 mM^-1^ s^-1^ which is comparable to GlgA2 at 3.4 mM^-1^ s^-1^. To further validate our assay method, we performed parallel ADP-Glc titration replacing the *Synechocystis* GlgA isoenzymes with *E. coli* GlgA (GlgA*^E.coli^*) under identical reaction conditions(**Figure 2B**). Notably, GlgA*^E.coli^* exhibited enzymatic activity approximately 10³-fold higher than *Synechocystis* GlgA1 and GlgA2, consistent with previous reports [16]. These results confirm the assay’s reliability across different GlgA enzymes, while highlighting the substantially slower catalytic rates of *Synechocystis* isoenzymes relative to *E. coli*.

**Figure 1.**
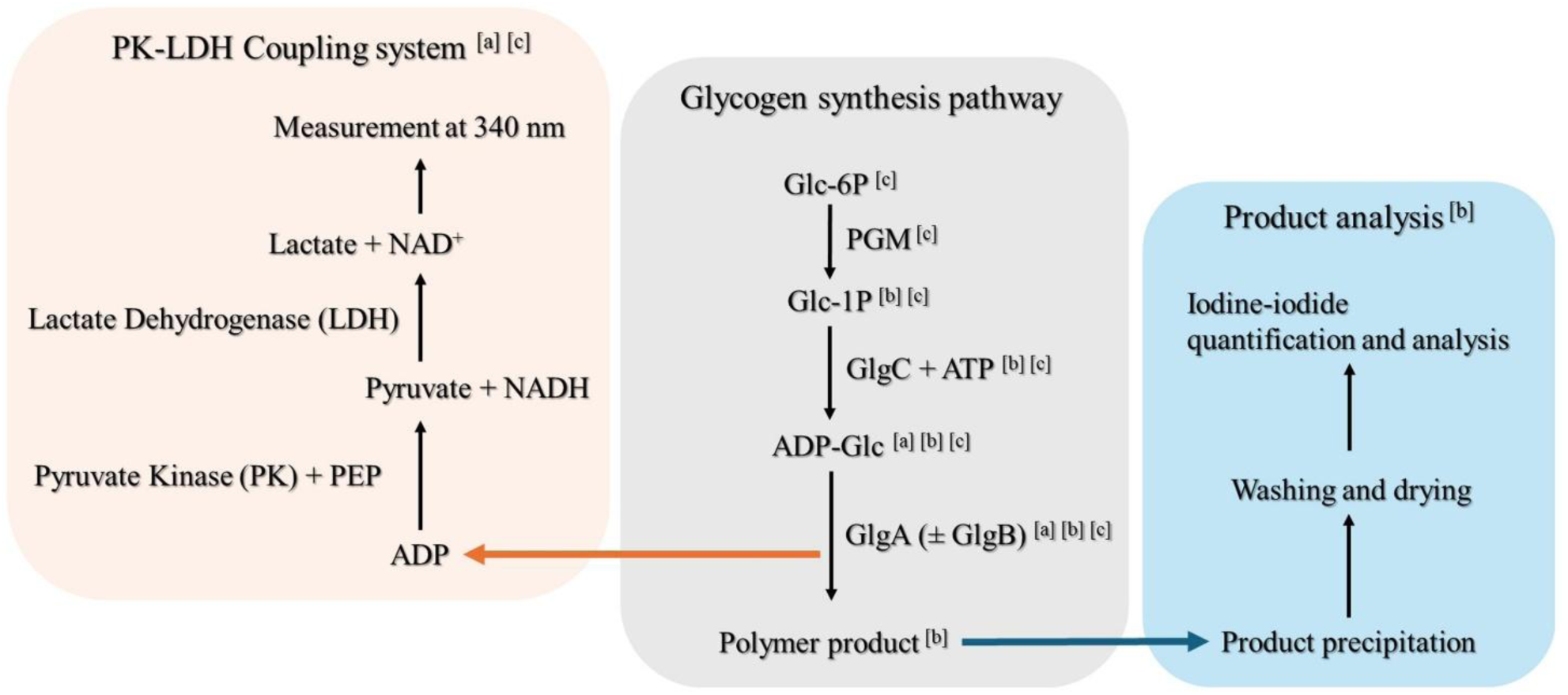
Schematic overview of the glycogen synthesis pathway in *Synechocystis*, and methods used in this study to assay different components of the pathway. Indicated are components [a] involved in Assay A [b] involved in Assay B [c] involved in Assay C. The details of each assay are given in their respective sections in the methods.

**Figure 2.**
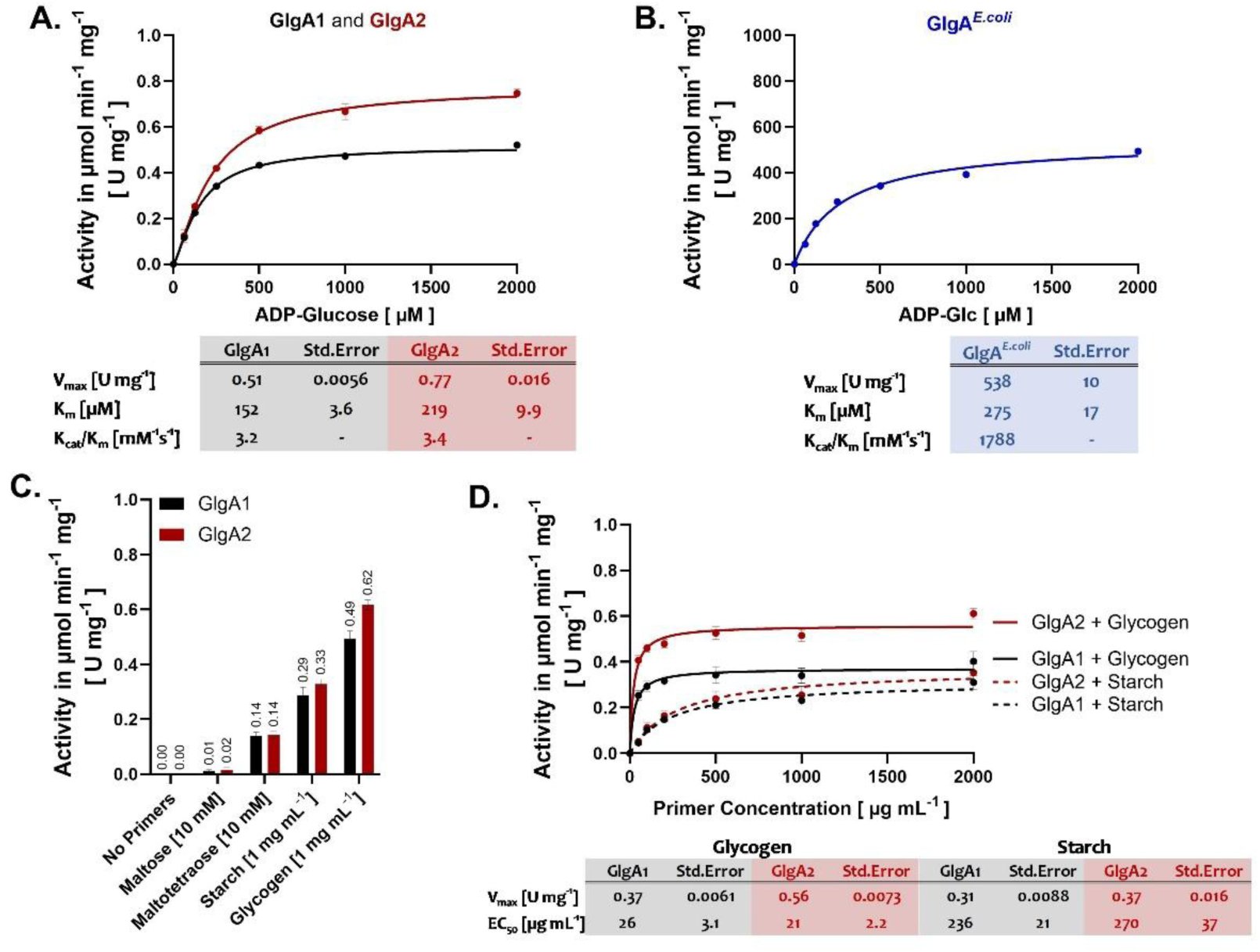
GlgA Kinetics assayed using the ADP/NADH Coupled Assay for glycogen synthesis Activity (Assay A). **(A)** ADP-Glc titration with 200 nM GlgA1 or GlgA2. 1 mg mL^-1^ glycogen was used as a primer. **(B)** ADP-Glc titration with 1 nM GlgA*^E.coli^* . 1 mg mL^-1^ glycogen was used as a primer. **(C)** Comparison between synthesis primers. The reaction was started with 1 mM ADP-Glc. **(D)** Titration curves for glycogen and starch. As above, the reaction was started with the addition of 1 mM ADP-Glc. Where applicable, each data point represents a mean of at least 3 replicates (n ≥ 3) with the SD shown as error bars.

### Primer architecture dictates GlgA activity profiles

Next, we assessed the impact of primer structure on glycogen synthesis by testing maltose and maltotetraose as alternative short and unbranched primers, alongside comparisons with soluble potato starch (hereafter starch) and bovine glycogen to evaluate the effect of primer branching. Both GlgA1 and GlgA2 utilized the shorter, less branched primers, albeit with reduced activity compared to glycogen (**Figure 2C**). No glycogen synthesis was observed in the absence of a primer. These results indicate that α-1,4-glycosidic bond formation by GlgA at non-reducing ends strictly requires priming, with a preference for longer, branched glycan primers.

To further evaluate primer-dependent modulation, starch and glycogen concentrations were titrated at near saturating ADP-Glc levels (1 mM) (**Figure 2D**). Both isoenzymes exhibited a rapid increase in activity with rising glycogen concentrations (EC₅₀: GlgA1 = 26 ± 3.1 µg mL⁻¹; GlgA2 = 21 ± 2.2 µg mL⁻¹), whereas responses to starch were significantly weaker (EC₅₀: GlgA1 = 236 ± 21 µg mL⁻¹; GlgA2 = 270 ± 37 µg mL⁻¹). Comparing maximal activities (V_max_), reactions using starch as a primer resulted in a reduction in GlgA1 activity by 16% and GlgA2 activity by 34% compared to glycogen. These data suggest that, despite comparable catalytic efficiencies, the isoenzymes differ in their primer interactions, potentially reflecting variant preferences for primer structure or branching.

### GlgB-driven branching preferentially enhances GlgA2 activity

Next, we introduced the branching enzyme GlgB into the glycogen synthesis reaction to promote primer branching and examined its effects on GlgA activity. Both GlgA isoenzymes exhibited noticeable activation with increasing GlgB concentrations up to around 20 nM GlgB or a molar ratio of one GlgB to ten GlgA monomers (**Figure 3A**). **Figure 3B** illustrates the net activation (activity normalized against a control without GlgB) of both GlgA isoenzymes with increasing concentrations of GlgB, using starch as a representative of an unbranched primer, demonstrating at least a twofold enhancement in their activity . However, the maximum activation of GlgA1 (increase of 0.17 ± 0.02 U mg⁻¹) was substantially lower than that of GlgA2 (increase of 0.37 ± 0.01 U mg⁻¹). Furthermore, GlgA1 required almost twice the amount of GlgB for half-maximal activation (EC₅₀: 7.8 ± 1.7 µM) compared to GlgA2 (EC₅₀: 4.4 ± 0.19 µM). These results suggest that GlgB activation of GlgA2 is slightly more efficient than that of GlgA1, which is further supported by the Hill coefficient for GlgA2 (n^H^: 1.9 ± 0.18) compared to GlgA1 (n^H^: 1.3 ± 0.22), indicating positive cooperativity between GlgB and GlgA2 (**Figure 3C**).

**Figure 3.**
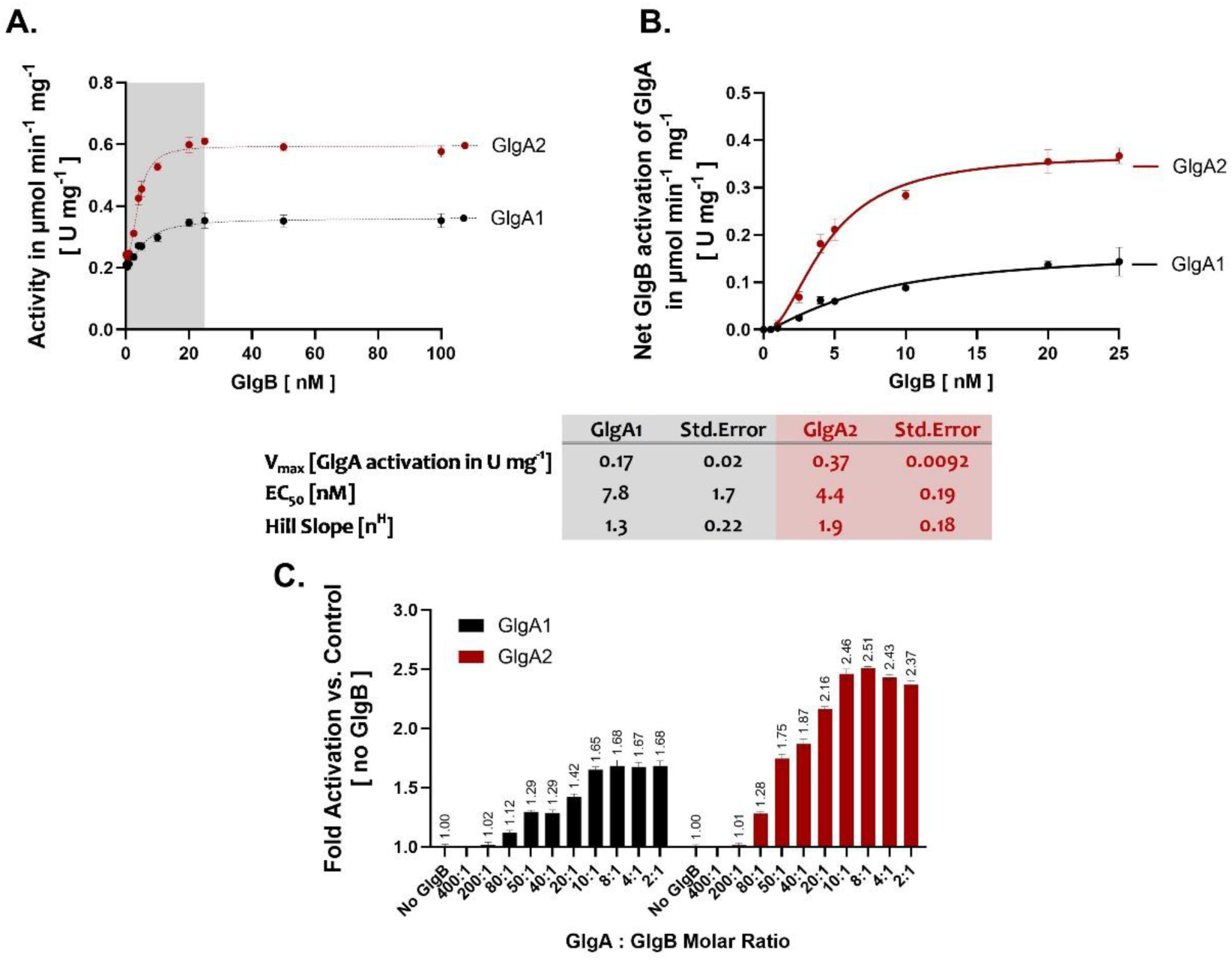
Activation of GlgA isoenzymes by GlgB. **(A)** GlgA enzymatic activity with GlgB. The assay was performed using increasing concentrations of GlgB with 200 µg mL⁻¹ soluble starch as the synthesis primer. Reactions were initiated by adding 1 mM ADP-Glc. The area shaded in grey is represented in panel **(B)**. **(B)** Net activation of both GlgA isoenzymes (200 nM GlgA) was assessed with increasing GlgB concentrations in the presence of 200 µg mL⁻¹ soluble starch as the synthesis primer. Reactions were initiated by adding 1 mM ADP-Glc. Net activity values on the y-axis represent GlgA activity after subtracting the baseline activity measured without GlgB from each corresponding data point on the x-axis. **(C)** The fold change in GlgA activity was calculated relative to a control reaction without GlgB. GlgB concentrations are expressed as molar ratios to 200 nM GlgA isoenzyme, with the optimal GlgA: GlgB ratio determined to be 10:1. Raw activity values are presented in panel **(A)**. Where applicable, each data point represents a mean of at least 3 replicates (n ≥ 3) with the SD shown as error bars.

To elucidate the modulatory effects of GlgB on GlgA isoenzyme activity, we tested increasing concentrations of starch in the presence of both enzymes to identify the primer concentration at which GlgB-mediated branching most effectively potentiates GlgA catalytic performance. In the presence of GlgB, the EC₅₀ for starch was approximately fourfold lower for GlgA2 than GlgA1, accompanied by a higher V_max_ for GlgA2 (**Figure 4A**). More strikingly, when GlgB cooperates with either GlgA isoenzyme, the EC₅₀ for starch decreased by at least 70-fold for GlgA1 and 330-fold for GlgA2 compared to reactions without GlgB (**Figure 4B**). Interestingly, the maximum activity of GlgA1 at saturating starch concentrations remained essentially unchanged by GlgB (V_max_ with GlgB: 0.30 ± 0.02 U mg⁻¹ vs. without GlgB: 0.31 ± 0.01 U mg⁻¹). In contrast, GlgA2 exhibited a 21.6% increase in maximum activity in the presence of GlgB (V_max_ with GlgB: 0.45 ± 0.01 U mg⁻¹ vs. without GlgB: 0.37 ± 0.02 U mg⁻¹). Moreover, the catalytic efficiencies (defined as K_cat_/Primer EC_50_) of both enzymes increased by at least two orders of magnitude in the presence of GlgB, highlighting the substantial impact of branching on enzyme performance (**Figure 4**). Notably, the catalytic efficiency of GlgA2 with GlgB was 6.4-fold higher than that of GlgA1, underscoring a strong dependence of GlgA2 activity on cooperative interactions with GlgB. Taken together, these findings indicate that GlgB significantly enhances the catalytic efficiency and substrate affinity of both GlgA isoenzymes, with a particularly pronounced effect on GlgA2, while GlgA1 activity at higher primer concentrations appears to be more influenced by the overall primer concentration rather than maximum activation by GlgB alone.

**Figure 4.**
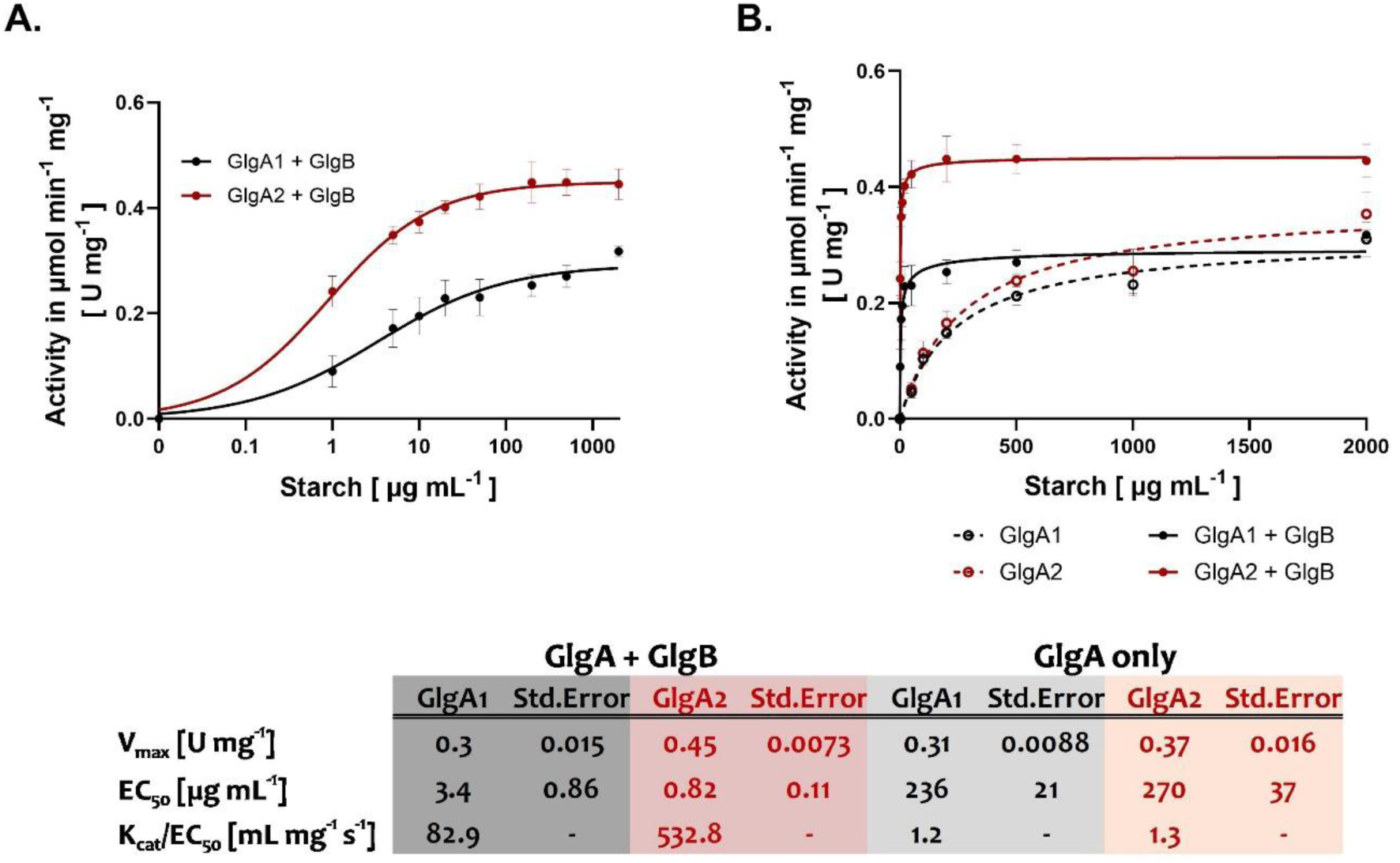
Effect of branching on GlgA primer sensitivity. **(A)** Starch titration in the presence of 20 nM GlgB and 200 nM GlgA (1:10 molar ratio of GlgB to GlgA) with 1 mM ADP-Glc, showing the effect of primer concentration on GlgA activity. **(B)** Overlay of starch titration curves with GlgB (solid lines), and corresponding starch titration data from Figure 2C (dashed lines)for direct comparison. Where applicable, each data point represents a mean of at least 3 replicates (n ≥ 3) with the SD shown as error bars.

### Comparative capacities identify GlgA as the rate-limiting step

To evaluate how coupling to the comparatively slower *Synechocystis* GlgA isoenzymes (as compared to GlgA*^E.coli^* affects GlgC, we compiled previously reported kinetics parameters for GlgC measured by malachite green (MG) Pi-release assay together with those for GlgA reported above. We compared the intrinsic catalytic capacities on a common flux scale (µM min⁻¹) to relate their activities directly. Using the MG assay-derived specific activity of GlgC (8.9 U mg⁻¹ at 4.98 nM GlgC tetramer) as a benchmark for supply, and the Assay A specific activities (V_max_) of GlgA1, GlgA2 and GlgA*^E. coli^* as measures of utilization capacity, we estimated the minimum GlgC concentration required to match GlgA demand under assay conditions (**Table 2**) [6]. For *Synechocystis*, our calculations indicate that matching the demand of 200 nM GlgA would require on the order of ∼3.2 nM GlgC for GlgA1 (≈62 GlgA1 monomers per GlgC tetramer) or ∼5 nM GlgC for GlgA2 (≈40 GlgA2 monomers per GlgC tetramer), whereas the highly active GlgA*^E. coli^* would require over 3.33 µM GlgC tetramer to match 200 nM of GlgA*^E.coli^*(≈0.06:1). Thus, *Synechocystis* GlgA remains the kinetic bottleneck over a wide range of realistic GlgA:GlgC ratios, while GlgA*^E. coli^*would be strongly supply-limited by GlgC.

### GlgA1 produces more-branched glycogen, whereas GlgA2 yields a more linear polymer

To study the products of the glycogen synthesis reaction in detail, we repurposed the Assay A system for preparative purposes by omitting the coupling system components for the spectrophotometric assays, namely LDH and NADH (**Figure 1**). To complete the *in vitro* glycogen-synthesis pathway, we introduced an upstream ADP-Glc production module by integrating GlgC at a stoichiometric ratio of 1:1 GlgA monomer to GlgC tetramer, and started the glycogen synthesis reaction via addition of Glc-1P, using starch as the synthesis primer (Assay B).

Products formed in the presence of GlgB were considered predominantly branched glycogen, whereas those produced in its absence were considered amylose-like linear glucans. Polymer synthesis products from Assay B were quantified by both iodine-iodide spectrophotometric staining and enzymatic determination (**Figure 5A**), yielding consistent results with no significant differences observed between the two quantification methods. **Figure 5B** presents the synthesis product yields for both GlgA isoenzymes with and without GlgB. From 1 mL reaction mixtures, GlgA2 in combination with GlgB produced 18.1% more glycogen (1.3 mg) than the GlgA1-GlgB reaction (1.1 mg). Similarly, in the absence of GlgB, GlgA2 synthesized 12.9% more amylose-like polymer (0.233 mg) compared to GlgA1 (0.206 mg). Notably, reactions lacking GlgB demonstrated a substantial decrease in product yield, with an 82.9% reduction for GlgA1 and 81.3% for GlgA2.

**Figure 5.**
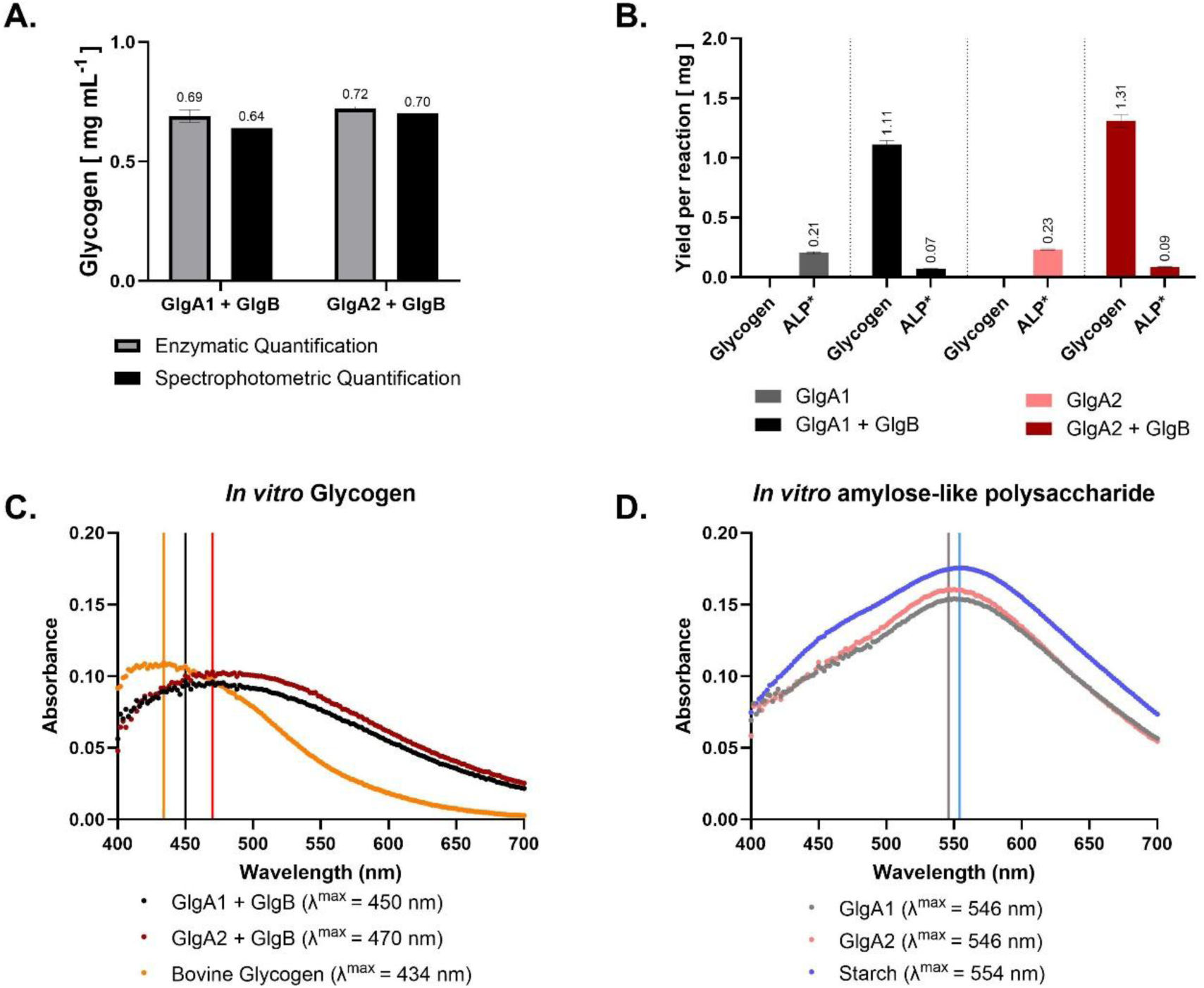
Characterisation of *in vitro* polymer products (Assay B) **(A)** Comparison of glycogen quantification methods. Pooled glycogen samples from Assay B replicates (n = 6) were adjusted to 1 mL volume. Glycogen concentrations were first measured using the iodine-iodide spectrophotometric assay and subsequently validated by an enzymatic rapid glycogen assay. Paired t-test analysis yielded a P value of 0.2578, indicating no significant difference (P > 0.05). **(B)** Yields of reaction products from Assay B, expressed as mg of product per 1 mL reaction volume. Each data point represents the mean of six replicates (n = 6), with standard deviation (SD) shown as error bars. Absorbance spectra between 400 and 700 nm of pooled synthesis products from Assay B. Equal volumes of replicate reactions from **(B)** were combined, and concentrations were adjusted to 0.5 mg mL⁻¹ for glycogen samples and 0.1 mg mL⁻¹ for amylose-like samples. Maximum absorbance values (λ^max^) were calculated using the Area Under the Curve (AUC) function in GraphPad Prism, referencing the highest peak identified, and are indicated by vertical lines on the x-axis. * Amylose-like Polysaccharides (ALP) **(C)** Absorbance spectra of glycogen samples from Assay B compared to bovine glycogen standard. **(D)** Absorbance spectra of amylose-like samples from Assay B compared to a starch standard.

Analysis of iodine-iodide absorbance spectra revealed subtle differences in polymer branching between products of the two GlgA isoenzymes in the presence of GlgB (**Figure 5C**). Glycogen synthesized by GlgA1 showed an absorbance maximum (λ^max^) of 450 nm with higher absorbance at lower wavelengths, indicative of a more branched structure, approximating the profile of highly branched bovine glycogen (λ^max^: 434 nm). Conversely, GlgA2-derived glycogen exhibited a λ^max^ of 470 nm, suggestive of comparatively fewer branches. The amylose-like polysaccharides generated in the absence of GlgB from both isoenzymes showed λ^max^ values near 546 nm (**Figure 5D**), characteristic of linear starch-like molecules, closely matching the profile of the starch standard λ^max^ of 554 nm. We further compared these *in vitro*-polymer profiles with glycogen isolated from *Synechocystis* mutants deficient in either GlgA1 (Δ*GlgA1*) or GlgA2 (Δ*GlgA2*) (**Figure 6A**). Glycogen from the Δ*GlgA2* mutant displayed a λ^max^ of 470 nm, resembling bovine glycogen (**Figure 6B**), whereas glycogen from Δ*GlgA1* had a higher λ^max^ of 516 nm, aligning more closely with starch (**Figure 6C**).

**Figure 6.**
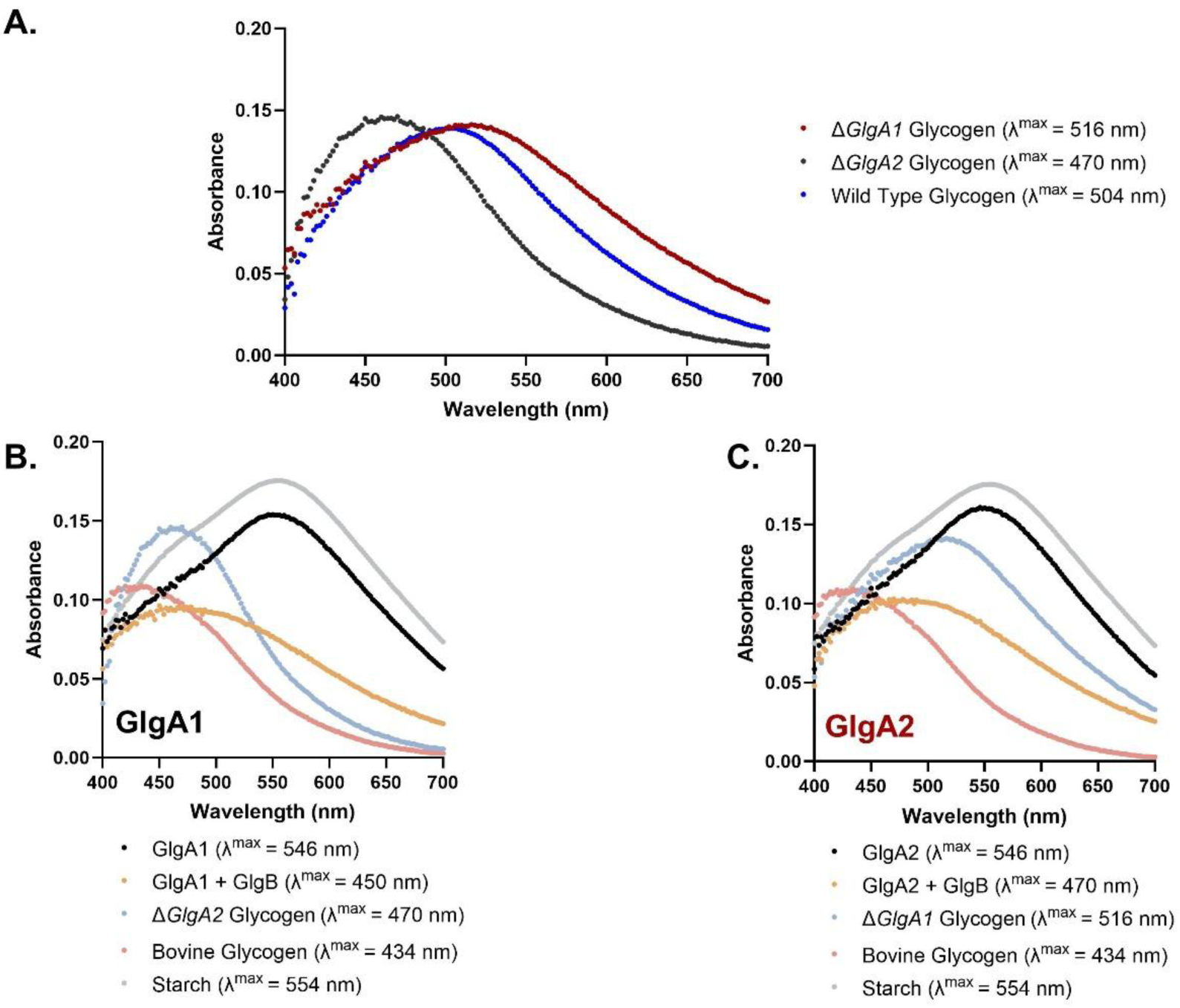
Comparison of *in vitro* polymer products with *in vivo* glycogen. Absorbance spectra between 400 and 700 nm of pooled synthesis products from Assay B were compared to glycogen extracted from *Synechocystis* wild type and GlgA-deficient mutant cultures. Equal volumes of replicate reactions from Assay B or isolated glycogen from *Synechocystis* cultures were combined, and concentrations were adjusted to 0.5 mg mL⁻¹ for glycogen samples and 0.1 mg mL⁻¹ for amylose-like samples. Maximum absorbance values (λ^max^) were calculated as previously described using the Area Under the Curve (AUC) function in GraphPad Prism. **(A)** Absorbance spectra of glycogen isolated from Δ*GlgA2*, Δ*GlgA1*, and wild type *Synechocystis* cultures. **(B)** Comparison of GlgA1-derived synthesis products with bovine glycogen, starch, and glycogen isolated from GlgA2-deficient (Δ*GlgA2*) *Synechocystis* mutants. **(C)** Comparison of GlgA2-derived synthesis products with starch, bovine glycogen, and glycogen isolated from GlgA1-deficient (Δ*GlgA1*) *Synechocystis* mutants.

Although neither mutant-derived glycogen spectrum perfectly matched their respective *in vitro* synthesized products, the glycogen produced *in vivo* by GlgA1 exhibited spectral properties (λ^max^ = 470 nm) closely resembling those of *in vitro* GlgA1-derived glycogen (λ^max^ = 450 nm). Both forms demonstrated a higher degree of branching compared to glycogen produced by GlgA2. *In vivo* synthesized glycogen by GlgA2 displayed starch-like characteristics with a notably higher λ^max^ of 516 nm, indicating less branching. This trend of reduced branching was also observed in glycogen synthesized *in vitro* by GlgA2, though less pronounced, as reflected by a λ^max^ of 470 nm. Wild-type glycogen showed intermediate properties, with a λ^max^ of 504 nm, positioning it between the two mutant profiles and recapitulating the distinct structural divergences between GlgA1- and GlgA2-derived glycogen *in vivo*. (**Figure 6A**).

### Modified Assay A enables quantification of PGM forward reaction kinetics

To further expand the utility of our assay system, we modified the core assay (Assay A) to examine PGM activity by coupling the PGM reaction to equimolar GlgC and GlgA*^E.coli^* (Assay C) (**Figure 1**). PGM catalyzes the reversible interconversion of Glc-1P and Glc-6P. *Synechocystis* bears two genes encoding putative PGM enzymes (*sll0726 and slr1334*); the sll0726-encoded enzyme was selected for this study, as it represents the primary PGM isoform in *Synechocystis* [17]. To ensure substrate turnover was not rate-limited, the coupling enzymes GlgC and GlgA*^E.coli^* were included in the reaction at 400 nM each, placing the GlgA:GlgC ratio at 1:1, far above the ideal 0.06:1 established previously (**Table 2**).

A titration of PGM was performed to identify the concentration range in which the measured activity reflected intrinsic enzyme catalysis (**Figure 7A**). The reaction rate increased linearly with PGM concentration up to the tested concentration of 6.13 nM, indicating that within this range, the observed activity corresponded directly to PGM turnover. Together with the determined dissociation constant (K_d_ : 6.4 ± 0.25 nM), these results indicate that enzyme concentrations withinthis range are optimal for reliable kinetic measurements under the applied conditions. Subsequently, a Glc-6P titration was conducted using 5 nM PGM. As shown in **Figure 7B**, the resulting kinetic parameters demonstrated that the assay effectively resolved PGM catalytic properties, yielding a V_max_ of 190 ± 6.6 U mg⁻¹ and a K_m_ of 1.52 ± 0.16 mM for Glc-6P.

**Figure 7.**
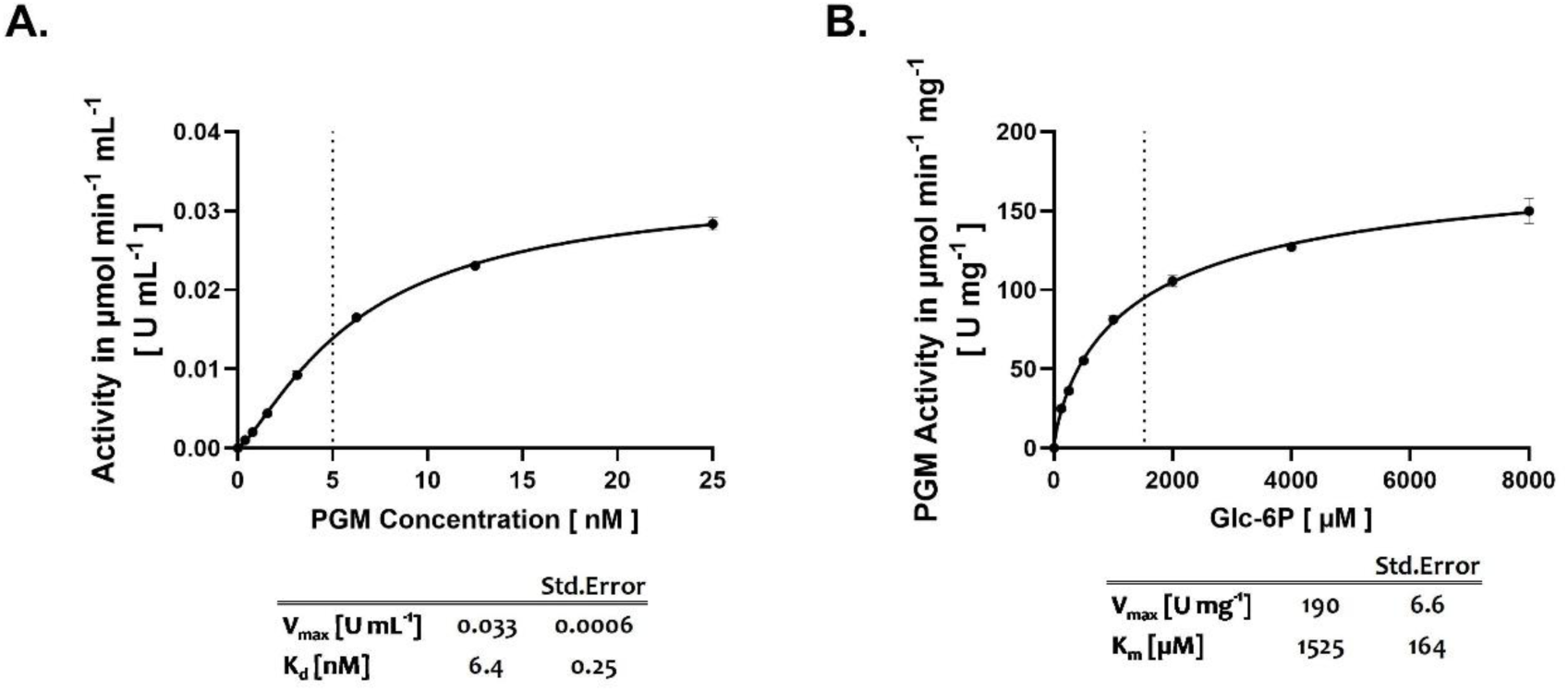
PGM forward reaction Assays (Assay C). **(A)** Titration of PGM at the indicated concentrations. Reactions were initiated by the addition of 1 mM Glc-6P. Activity values (U mL⁻¹) on the y-axis represent the overall enzymatic activity of the coupled pathway, including Glc-1P formation from Glc-6P, subsequent ADP-Glc synthesis, and glycogen polymerization. The dotted line on the x-axis denotes the standard PGM concentration used in subsequent assays. **(B)** Titration of Glc-6P in the presence of 5 nM PGM. The K_m_ value is indicated by the dotted line on the x-axis. Where applicable, each data point represents a mean of at least 3 replicates (n ≥ 3) with the SD shown as error bars.

## Discussion

We established an *in vitro* model of glycogen synthesis pathway from *Synechocystis* (**Figure 1**). This model elucidates critical factors controlling glycogen synthesis, including primer selectivity and the modulatory role of GlgB. Additionally, we demonstrated production of glycogen *in vitro* and characterized structural differences between products synthesized by GlgA1 and GlgA2.

### Primer architecture and GlgB cooperation partition elongation labour between GlgA isoenzymes

The kinetic analyses demonstrate that GlgA1 and GlgA2 isoenzymes possess broadly similar catalytic efficiencies despite subtle differences in their kinetic parameters, implying differential optimization potentially aligned with distinct cellular or metabolic contexts. Both isoenzymes can utilize short, linear polysaccharide primers, though less efficiently than branched glycogen, indicating some flexibility in primer selectivity but also emphasizing that primer structure notably influences enzymatic activity. The strict primer requirement, evidenced by the absence of glycogen synthesis without a primer, underscores a fundamental primer-dependent mechanism and suggests additional, yet unidentified factors may facilitate glycogen initiation in *Synechocystis* beyond GlgC and GlgB (**Figure 2C**) [18]. Furthermore, the distinct responses of GlgA1 and GlgA2 to starch and glycogen highlight differential primer interactions. Significantly higher concentrations of linear starch were needed to reach activities comparable to those with branched glycogen, indicating that branched primers with more readily accessible non-reducing ends are essential for optimal GlgA binding and activation (**Figure 2C**, **D**). This highlights the role of GlgB in glycogen synthesis, where the inclusion of GlgB markedly enhanced the catalytic activities of both GlgA isoenzymes, albeit to different extents (**Figure 3**). GlgA2, which synthesizes longer glucan chains, is suggested to be particularly susceptible to steric hindrance within the growing glycogen network [5,10,19]. The branching activity of GlgB alleviates these steric constraints by creating new branch points, thereby enabling GlgA2 to extend even longer chains more efficiently [10]. Consequently, GlgA2 exhibited a substantially stronger and more persistent activation in the presence of GlgB, indicative of a highly efficient functional cooperativity between the two enzymes (**Figure 3B**). This pronounced synergy likely reflects a close coordination of elongation and branching reactions that supports continuous and effective glucan synthesis. In contrast, GlgA1 performs a more dispersive elongation, resulting in the synthesis of glycogen that is inherently more branched than that produced by GlgA2 [5,10]. Accordingly, GlgA1 activity was observed to be increased only up to a defined level and then plateaued in the presence of GlgB, suggesting that the additional branching generated by GlgB does not further stimulate catalysis beyond a certain limit (**Figure 3**). Additionally, at elevated GlgB concentrations, a slight decrease in GlgA2 activity was observed (**Figure 3C**), an effect absent in GlgA1. This pattern suggests that GlgA2 may be particularly sensitive to steric effects arising from excessive enzyme crowding or suboptimal access to elongation sites despite branching’s general role in relieving steric hindrances. Taken together, these observations highlight that GlgA2 engages in significantly more efficient and finely tuned cooperation with GlgB than GlgA1. The iodine-iodide spectra of products formed with GlgB further reinforce that GlgA1 glycogen is more branched than GlgA2 glycogen despite similar primer usage [5,18].

This aligns the observations from our enzyme assays with the physicochemical differences observed between glycogen synthesized by the two GlgA isoenzymes: glycogen produced by GlgA1, both *in vivo* and *in vitro*, is notably more branched than that formed by GlgA2 (**Figure 5C**, **Figure 6A**). In the absence of GlgB, both isoenzymes generate amylose-like, largely linear polymers with largely indistinguishable iodine-iodide spectral profiles, likely reflecting some degree of synthesis inhibition or a drastic reduction in GlgA activity to negligible levels over longer durations of sustained synthesis in the absence of branching (**Figure 5D**). Taken together with the markedly lower yields of amylose-like polysaccharides in reactions without GlgB, it is reasonable to conclude that branching is also essential to relieve steric hindrances that occur during sustained glycogen synthesis (**Figure 5B**, **D**) [19]. These findings demonstrate that the role of GlgB extends beyond primer branching —it also plays a key role in glycogen structure reorganisation for efficient GlgA activity and allows for efficient sustained glycogen synthesis.

The structural distinctions in glycogen produced by each GlgA isoenzyme, in addition to the characteristics of the isoenzymes themselves may reflect specialized physiological roles. For example, the short, highly branched glycogen synthesized by GlgA1 could confer survival advantages under environmental stresses, as observed in various bacteria where such glycogen is degraded more slowly, supporting reduced metabolic rates and prolonged viability, given that the shorter branches are more easily accessible to glycogen degradation enzymes [13]. In *Synechocystis*, the dependence of resuscitation from nitrogen starvation on GlgA1 supports the notion that GlgA1-derived glycogen contributes to adaptability and survival, whereas GlgA2-generated glycogen primarily could serve as a carbon storage molecule during conditions of high carbon flux [14,18,20]. Given that sustained glycogen synthesis occurs under such conditions, it is reasonable to assume that the reorganisation of the growing glycogen granule by GlgB is especially crucial to support optimal GlgA2 activity. Since wild-type glycogen exhibits characteristics intermediate between those of GlgA1- and GlgA2-derived glycogen, it is reasonable to conclude that both isoenzymes cooperatively contribute to glycogen synthesis during normal growth, balancing metabolic flexibility with carbon storage needs (**Figure 6A**) [10].

### Divergent glycogen synthesis control in phototrophs and heterotrophs

Oxygenic phototrophs—including *Synechocystis*—allocate primary control upstream at GlgC, which is activated by 3-PGA and inhibited by Pi [6,8,9,21], and diversify elongation via two GlgA isoenzymes with distinct operational biases [5,18]. Additionally, given the roughly comparable *in vivo* expression levels of GlgA monomers and GlgC tetramers during normal heterotrophic growth, our observations also suggest an additional layer of intrinsic regulation of glycogen synthesis via enzyme stoichiometry [20,22,23]. Our *in vitro* reconstitutions support this proposed architecture in that: (i) GlgB sharply potentiates both GlgA isoenzymes under low-primer conditions but disproportionately enhances GlgA2, (ii) product spectra indicate that GlgA1 yields more highly branched glycogen (lower λ^max^) than GlgA2, and (iii) at a practical and biologically relevant 1:1 GlgC tetramer : GlgA monomer ratio, pathway flux is GlgA-limited due to the lower intrinsic activity of the *Synechocystis* GlgA isoenzymes. In sum, primer/branching governance and modest GlgA catalysis—rather than maximal elongation throughput—determine the glycogen synthesis flux and polymer architecture in *Synechocystis*.

By contrast, in heterotrophic bacteria such as *E. coli*, glycogen synthesis is configured for rapid accumulation. GlgA*^E.coli^* has high catalytic capacity *in vitro* [16,24], and the GlgA–GlgB pair sustains appreciable elongation even when exogenous primer is scarce—under “unprimed” conditions, GlgB (further facilitated by citrate) supports ∼30% of the GlgA*^E.coli^* V_max_ otherwise achieved with saturating glycogen primer concentrations [25,26]. *In vivo*, the maltose/maltodextrin network (MalQ/MalP and associated transporters) continually supplies maltooligosaccharide acceptors, minimizing primer limitation and amplifying the impact of a fast GlgA [27,28]. These features indicate an evolutionary reason why *E. coli* GlgC (GlgC*^E.coli^*) exhibits substantially higher intrinsic activity than the *Synechocystis* isoenzymes, and why GlgA*^E.coli^* activity exceeds GlgC*^E.coli^* by several fold [6,24,29].

A further distinction concerns initiation and primer economy. Bacteria do not employ a glycogenin-type protein primer; in some species, glycogen synthesis can initiate *de novo*, and primer pools arise via enzyme-mediated interconversion of maltooligosaccharides [30]. This helps explain why *E. coli* can partly circumvent primer scarcity through GlgB-assisted “unprimed” synthesis and MalQ/MalP-driven primer provision [26–28]. On the other hand, *Synechocystis* lacking a dedicated maltose circuit, relies more on primer economy and GlgB cooperation to maintain throughput. A similar control logic of GlgA-GlgB coordinated elongation–branching is also documented in higher plants, where multiple starch-synthase and branching-enzyme isoforms assemble into catalytically competent complexes [31,32]. Additionally, similar GlgA-GlgC catalytic arrangements—higher intrinsic GlgC activity relative to starch/glycogen synthases—were also observed in various higher plants, hinting that these regulatory strategies might be common to photoautotrophs [1,33–37]. Viewed together, these considerations provide a plausible mechanistic rationale: *Synechocystis* GlgA operates in a measured kinetic regime that facilitates efficient GlgB-mediated branch insertion under primer-limited conditions, rather than being tuned for maximal elongation velocity per se, contrasting with the primer-rich, high-throughput configuration of *E. coli*.

### The coupled assay (Assay B) applications go beyond modelling glycogen synthesis

The coupled assay system developed in this study demonstrates considerable flexibility and can be readily adapted to address a broad range of biochemical questions beyond glycogen synthesis itself. Its modular design allows precise modification of reaction components to target different catalytic processes or synthetic outcomes. For example, scaling the assay for preparative purposes (Assay B, **Figure 5**) highlights its utility for large-scale *in vitro* glycogen synthesis, providing new perspectives on glucan structure formation and enzyme cooperation. Beyond analytical use, such adaptations may also serve practical applications, such as generating defined *Synechocystis* glycogen for downstream enzymatic studies—most notably of glycogen phosphorylase—without requiring extensive biomass cultivation.

Apart from applications directly related to glycogen synthesis, the adaptation applied in Assay C (**Figure 7**) establishes a powerful approach for investigating phosphoglucomutase (PGM) activity and its regulation. While the kinetics of the reverse PGM reaction (Glc-1P → Glc-6P) have been thoroughly examined, the forward conversion (Glc-6P → Glc-1P) has received far less attention [17,38] . By coupling GlgC and GlgA*^E.coli^*, our system enables reliable quantification of this underexplored reaction and could be easily expanded to screen for potential metabolic regulators influencing PGM activity. Collectively, these applications underscore the assay’s versatility as a platform for probing diverse aspects of carbohydrate metabolism and enzymatic regulation in a controlled *in vitro* context.

## Conclusion

Our reconstruction of the *Synechocystis* glycogen biosynthesis pathway *in vitro* resolves glycogen synthesis into a hierarchical control scheme and a context-dependent division of labour between GlgA isoenzymes. Upstream, GlgC governs precursor supply: its substrate affinity and allosteric effectors tune ADP-glucose availability and thereby gate entry into the pathway [6]. Downstream, GlgA imposes the principal flux limitation. Within this framework, primer architecture and GlgB cooperation partition elongation tasks between GlgA1 and GlgA2—branching density selectively accelerates GlgA2-mediated chain extension, whereas GlgA1 favours shorter, more-branched products—linking flux control to product fine structure. Finally, the coupled assay we established offers a modular platform to quantify control coefficients and to generalise these principles to other nucleotide-sugar polymer pathways, such as resolving PGM forward kinetics.

## Methods

### Culture and growth conditions of E. coli

Unless otherwise specified, the cultivation of all strains was performed as previously reported [6]. All strains used in this study are listed in Table 1.

**Table 1:**
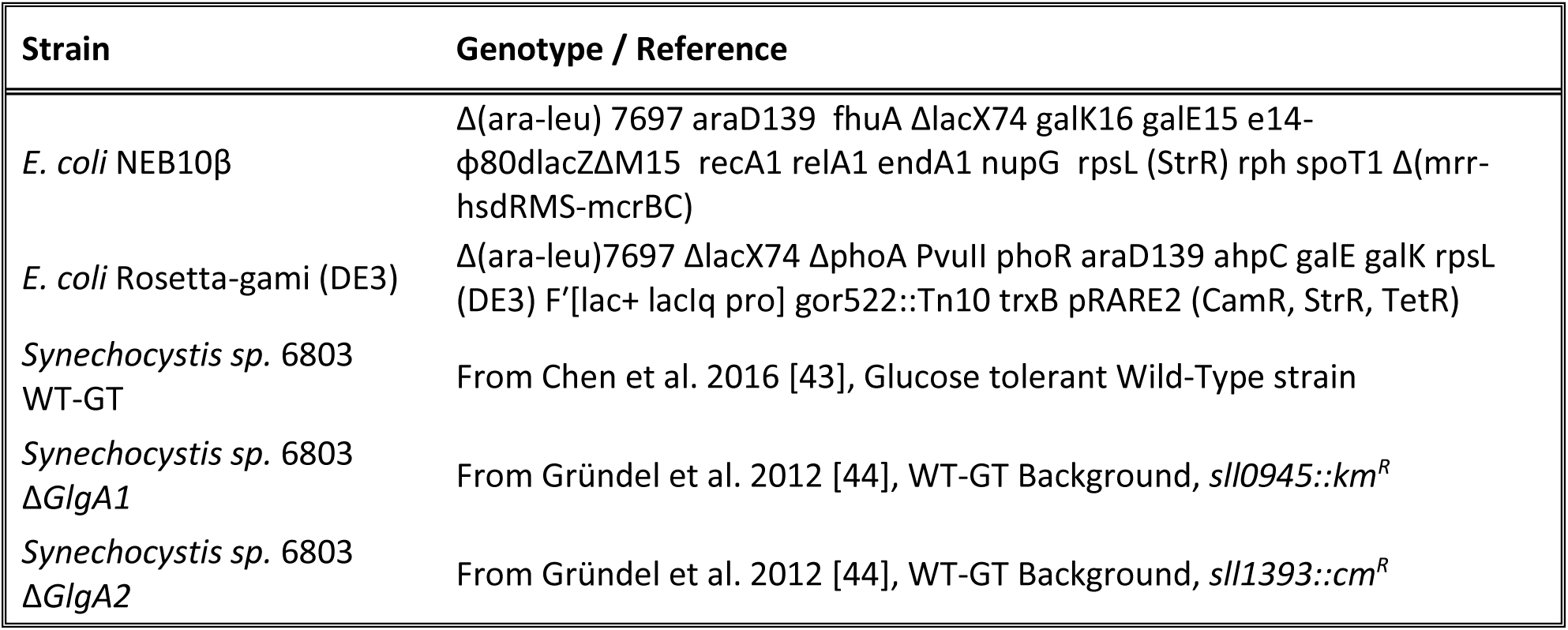
*E. coli* and *Synechocystis sp*. PCC6803 strains used in this work.

| Strain | Genotype / Reference |
| --- | --- |
| *E. coli* NEB10β | $\Delta(\text{ara-leu})$ 7697 $\text{araD139}$ $\text{fhuA}$ $\Delta\text{lacX74}$ $\text{galK16}$ $\text{galE15}$ $\text{e14-}\phi 80\text{dlacZ}\Delta\text{M15}$ $\text{recA1}$ $\text{relA1}$ $\text{endA1}$ $\text{nupG}$ $\text{rpsL}$ (StrR) $\text{rph}$ $\text{spoT1}$ $\Delta(\text{mrr-hsdRMS-mcrBC})$ |
| *E. coli* Rosetta-gami (DE3) | $\Delta(\text{ara-leu})$7697 $\Delta\text{lacX74}$ $\Delta\text{phoA}$ $\text{PvuII}$ $\text{phoR}$ $\text{araD139}$ $\text{ahpC}$ $\text{galE}$ $\text{galK}$ $\text{rpsL}$ (DE3) $\text{F}'[\text{lac+ lacIq pro}]$ $\text{gor522::Tn10}$ $\text{trxB}$ $\text{pRARE2}$ (CamR, StrR, TetR) |
| *Synechocystis* sp. 6803 WT-GT | From Chen et al. 2016 [43], Glucose tolerant Wild-Type strain |
| *Synechocystis* sp. 6803 $\Delta\text{GlgA1}$ | From Gründel et al. 2012 [44], WT-GT Background, $\text{sll0945::km}^R$ |
| *Synechocystis* sp. 6803 $\Delta\text{GlgA2}$ | From Gründel et al. 2012 [44], WT-GT Background, $\text{sll1393::cm}^R$ |

### Culture and growth conditions of Synechocystis sp. PCC6803

The cultivation of all cyanobacterial strains was performed as previously described [14]. All strains used in this study are listed in Table 1.

### Expression and purification of recombinant His-Tagged glycogen synthesis enzymes

The assembly of the expression constructs of N-terminal His-tagged GlgA1 and GlgC proteins was performed previously as described [6].

The GlgA2 gene was amplified from Synechocystis sp. PCC 6803 genomic DNA using the following primers; forward: *5’-GCCTGGTGCCGCGCGGCAGCATGTACATCGTTCAAATTGCCTCAGAATGCGC-3’*and reverse: *5’-TGTCGACGGAGCTCGAATTCTTAAGCCCGGATGTATTCGTAGGCTTCCACA-3’.*The expression plasmid pET28a was similarly linearized and amplified using the following primer pair; forward: *5’-GCAATTTGAACGATGTACATGCTGCCGCGCGGCACC-3’* and reverse: *5’-ACGAATACATCCGGGCTTAAGAATTCGAGCTCCGTCGACAAGCTTGCGGC-3’*.

Similarly, the GlgA gene from *E. coli* (GlgA*^E.coli^*) was amplified from genomic DNA using the following primers; forward: 5’-GCCTGGTGCCGCGCGGCAGCATGCAGGTTTTACATGTATGTTCAGAGATGTTCCCG-3’ and reverse: *5’-TGTCGACGGAGCTCGAATTCCTATTTCAAGCGATAGTAAAGCTCACGGTACGACTTCG-3’.* The expression plasmid pET28a was similarly linearized and amplified using the following primer pair; forward: *5’-TTTACTATCGCTTGAAATAGGAATTCGAGCTCCGTCGACAAGCTTGC-3’* and reverse: *5’-CATACATGTAAAACCTGCATGCTGCCGCGCGGCACC-3’*.

The subsequent assembly of the pET28a expression constructs for GlgA2 and GlgA*^E.coli^* was carried out in the same manner as for GlgA1 and GlgC. The subsequent induction and expression of recombinant proteins were performed using IMAC as previously described [6]. Cell lysis was carried out in lysis buffer (20 mM Tris-HCl pH 7.8, 50 mM NaCl, 5 mM MgCl_2_, 40 mM imidazole) supplemented with CellLytic buffer, Benzonase® (both from Merck, Darmstadt, Germany), lysozyme and one tablet of cOmplete™ protein inhibitor cocktail (Roche, Mannheim, Germany). The lysate was clarified via ultracentrifugation at 16000 x g, 30 minutes at 4°C and filtered using a 0.22 µm syringe filter prior to column application.

The recombinant proteins were purified via immobilised metal affinity chromatography (IMAC) followed by a polishing step using anion exchange to yield proteins of a high purity. All purifications steps were performed using the ÄKTA purifier chromatography system (Cytiva, Freiburg, Germany). IMAC was performed with Ni-NTA HisTrap columns (Cytiva, Freiburg, Germany) according to the manufacturer’s directions. The His-Tagged proteins were eluted with a 30 CV gradient of 40 mM to 500 mM imidazole in a buffer containing 20 mM Tris-HCl pH 7.8, 500 mM NaCl. Peak fractions were identified based on their UV 280 nm signal and pooled.

Buffer exchange of the pooled fractions was performed with the Amicon® Ultra Centrifugal Filter, 30 kDa MWCO (Merck, Darmstadt, Germany) with two washes with IEX buffer (20 mM Tris-HCl pH 7.8) prior to anion exchange with HiTrap Q High Performance columns (Cytiva, Freiburg, Germany) according to the manufacturer’s directions. The proteins were eluted with a 30 CV gradient of 0 to 750 mM NaCl in IEX buffer (20 mM Tris-HCl pH 7.8). As before, fractions from the largest peak were collected and buffer exchange was performed with two washes of concentration buffer (20 mM Tris-HCl pH 7.8, 150 mM KCl, 1 mM EDTA) followed by exchange into storage buffer (20 mM Tris-HCl pH 7.8, 150 mM KCl, 1 mM EDTA, 50%_(v/v)_ glycerol).

### Expression and purification of recombinant Strep-Tagged proteins

The construct for recombinant C-terminal StrepII-tagged GlgB was purified as previously described [39]. PGM was purified as previously described [17].

### ADP/NADH Coupled Assay for GS Activity (Assay A)

Glycogen synthase activity was assayed as previously described using a modified spectrophotometric method from that presented by Wayllace et al. [6,40]. The following assay parameters were changed to suit the requirements for this study. The components of the assay in a total volume of 100 µL per reaction were; assay buffer (50 mM HEPES-NaOH, pH 8.0, 12 mM MgCl_2_) with 10 U mL^-1^ Lactate Dehydrogenase (LDH), 9 U mL^-1^ Pyruvate Kinase (PK), 1 mM Phosphoenolpyruvate (PEP), 0.4 mM NADH, 1 mg mL^-1^ Bovine Glycogen and 200 nM GS (GlgA1 or GlgA2). The reaction was assayed in 96-well clear bottom plates at 30°C for variable durations, depending on the experimental design. Absorbance was measured using Tecan Spark 10M (Tecan, Männedorf, Switzerland) with readings taken every minute at 340 nm. Each standard reaction was started with the addition of the substrate ADP-Glc.

### Glycogen synthesis and precipitation (Assay B)

The synthesis of glycogen was performed in a similar manner to that of assay A with several modifications. The components of the assay in a total volume of 5 mL per reaction were:; assay buffer (50 mM HEPES-NaOH, pH 8.0, 12 mM MgCl_2_) with 4 mM ATP, 4 mM 3-PGA, 4 mM Glc-1P and 200 µg mL^-1^ starch as a primer. 9 U mL^-1^ PK and 2 mM PEP were also added to maintain ATP concentrations for optimal GlgC activity. Glycogen synthesis enzymes were added in a concentration ratio of 200 nM monomeric GS to 200 nM tetrameric GlgC, with and without 20 nM of GlgB. The reaction was left to run overnight (∼20 hours) at 30°C with gentle agitation on a shaker.

The next day, all samples were heated at 90°C for 10 minutes to inactivate the protein components of the reaction, followed by centrifugation at 17000 x g, 4°C for 30 minutes to separate the insoluble fraction of the reaction. The supernatant was carefully decanted and mixed with 4 volumes of ice-cold absolute ethanol before being stored overnight at -20°C to precipitate the reaction products. The precipitate was then pelleted via centrifugation at 17000 x g, 4°C for 15 minutes and washed twice with 70% ethanol and 100% ethanol before being left to dry completely in a 60°C oven. The dried products were resuspended in distilled water and an additional filtration step was performed using Amicon® Ultra Centrifugal Filter, 3 kDa MWCO (Merck, Darmstadt, Germany) with four washes of distilled water to remove soluble contaminants. The remaining concentrate was resuspended in distilled water for further analysis.

### PGM Forward Reaction Assay (Assay C)

The PGM forward reaction assay was adapted from Assay A and the coupled assay described in our previous report [6]. Each reaction (100 µL total volume) contained assay buffer (50 mM HEPES-NaOH, pH 8.0, 12 mM MgCl_2_), 40 µM Glc-1,6-phosphate, 2 mM ATP, 2 mM 3-PGA, 1 mM PEP, 0.4 mM NADH, and 1 mg mL⁻¹ bovine glycogen as primer, together with the coupling enzymes LDH and PK as specified above. Glycogen synthesis was catalyzed by 400 nM monomeric GlgA*^E.coli^* and 400 nM tetrameric GlgC in the presence of PGM. Reactions were performed in 96-well clear-bottom plates as described previously and initiated by the addition of Glc-6P.

### GlgC tetramer concentration ratios required to match GlgA activity under assay conditions

We calculated the theoretical GlgA–GlgC stoichiometries on a flux basis using the intrinsic specific activities from Assay A (GlgA1, GlgA2, GlgA*^E. coli^* with ADP-Glc) and the previously reported intrinsic GlgC activity from the MG assay [6]

**Table 2**. For a given enzyme, the flux was expressed in µM min⁻¹per nanomolar enzyme, as 𝐽 = 𝑉_𝑚𝑎𝑥_ ·𝑀𝑊 · 10^−6^, where V_max_ was expressed in U mg⁻¹, MW is the molecular weight of GlgA monomers or GlgC tetramers in g mol⁻¹, and 1 U = 1 µmol min⁻¹. For GlgC, the supply capacity was computed as *J_C_*, while for each GlgA isoform the consumption capacity per nM was computed as *J_A_* . The GlgA : GlgC stoichiometric ratio required to match the GlgA demand was then obtained as 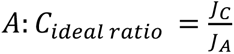 and the corresponding ideal concentration of GlgC required was calculated as 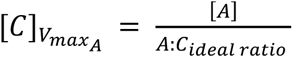. These calculations were used to compare the intrinsic supply capacity of GlgC with the utilization capacities of GlgA1, GlgA2 and GlgA*^E. coli^*, and to interpret the minimum GlgC concentration required for a stable supply of ADP-Glc to GlgA under assay conditions. **Table 2** summarizes all relevant parameters for this calculation.

**Table 2:** Experimental assay parameters and enzyme concentrations with molecular weights.

| Enzyme | Molecular Weight<br>[g mol<sup>-1</sup>] | Substrate | Spec. Activity<br>[U mg<sup>-1</sup>] | Flux Capacity<br>[$\mu\text{M min}^{-1}$ per nM] | K<sub>m</sub> [$\mu\text{M}$] | Optimal GlgA: GlgC ratio |
| --- | --- | --- | --- | --- | --- | --- |
| GlgC Tetramer | 200856.64 | Glc-1P | 8.90 | 1.79 | 240 ± 50 | — |
| GlgA1 | 56372.5 | ADP-Glc | 0.51 | 2.86 $\times 10^{-2}$ | 152 ± 3.6 | 62:1 |
| GlgA2 | 58251.0 | ADP-Glc | 0.77 | 4.49 $\times 10^{-2}$ | 219 ± 9.9 | 40:1 |
| GlgA<sup>*E.coli*</sup> | 54848.6 | ADP-Glc | 538 | 2.95 $\times 10^1$ | 275 ± 17 | 0.06:1 |

### Glycogen purification from Synechocystis cultures

Exponential *Synechocystis* cultures of 300 mL at ∼0.8 OD_750_ were harvested via centrifugation and washed twice with 30%_(w/v)_ KOH. The washed cell pellets were resuspended in 30%_(w/v)_ KOH and heated at 85°C for 30 minutes for cell lysis. After lysis, the precipitation of glycogen was carried out as previously described; the lysate was centrifuged at 17000 x g, 4°C for 30 minutes to separate cell debris. The supernatant was carefully decanted and mixed with 4 volumes of ice-cold absolute ethanol before being stored overnight at -20°C to precipitate glycogen. The precipitate was then pelleted via centrifugation at 17000 x g, 4°C for 15 minutes and washed twice with 70% ethanol and 100% ethanol before being left to dry completely in a 60°C oven. The dried products were resuspended in distilled water and an additional filtration step was performed using Amicon® Ultra Centrifugal Filter, 3 kDa MWCO (Merck, Darmstadt, Germany) with four washes of distilled water to remove soluble contaminants. The remaining concentrate was resuspended in distilled water for further analysis.

### GS reaction product quantification and analysis (Spectrophotometric quantification: Iodine-Iodide assay)

The quantification of the *in vitro* reaction products was performed by mixing the reaction products with 70 µL of iodine solution (2 mM I_2_, 30 mM KI) in a total volume of 150 µL. The reaction products were compared to glycogen or starch standard solutions in the concentration range of 0 to 2 mg mL^-1^ based on their absorbances at wavelengths of 490 nm or 560 nm for glycogen and starch respectively. For the measurement of the reaction product absorbance spectra, the concentration of the reaction products was adjusted to 0.5 mg mL^-1^, and the absorbance was measured from 400 nm to 700 nm. Measurements were performed using the Tecan Spark 10M (Tecan, Männedorf, Switzerland).

### Fast glycogen measurement assay (Enzymatic quantification)

The enzymatic determination of glycogen was performed with the method presented by Vidal et al. with some minor modifications, and was used to validate our Iodine-Iodide assay described above [41]. The total assay volume was modified to 100 µL, consisting of 30 µL enzymatic mix and 70 µL of glycogen sample from Assay B. The concentration of the glycogen sample was previously determined using the Iodine-Iodide assay and was used for comparisons. All measurements were done using the Tecan Spark 10M (Tecan, Männedorf, Switzerland).

### Figures, Bioinformatics, and Analyses

All data were analysed and visualized using GraphPad Prism 10 (GraphPad, CA, USA). Primer and expression construct designs were carried out in silico using SnapGene (GSL Biotech, CA, USA).

## Author contributions

KL: designed and carried out experiments, and wrote paper. DB: Assistance with Assay C. SD & KF: conceptualization, supervision and paper writing. All authors contributed at varying stages in the editing and review of the paper.

## Acknowledgements

We thank Dr. Christophe Colleoni at the University of Lille for his advice regarding the analysis of our synthesis products. This work was supported by the DFG funded research consortium FOR2816 “The Autotrophy-Heterotrophy Switch in Cyanobacteria: Coherent Decision-Making at Multiple Regulatory Layers”, research grant Fo195/16-2. We also acknowledge infrastructural Cluster of Excellence EXC 2124 (Controlling Microbes to Fight Infections, CMFI, grant 390838134) at the Eberhard Karls Universität Tübingen.

## Data availability

Enzymatic data and all figures presented in this study have been deposited here (DOI: 10.15490/fairdomhub.1.study.1417.2) on the data and model management platform FAIRDOMHub [42].

## Abbreviations

GlgA: Glycogen Synthase
GlgB: Glycogen Branching enzyme
GlgC: Glucose-1-phosphate adenylyltransferase
Glc-1P: Glucose-1-phosphate
Glc-6P: Glucose-6-phosphate
ADP-Glc: ADP-Glucose
Pi: Inorganic orthophosphate
Ppi: Pyrophosphate
3-PGA: 3-phosphoglycerate
LDH: Lactate dehydrogenase
PK: Pyruvate kinase
PEP: Phosphoenolpyruvate
MG: Malachite Green (Assay)
PGM: Phosphoglucomutase
*Synechocystis*: *Synechocystis* sp. PCC 6803

## References

1 McFadden GI (2001) Chloroplast Origin and Integration1. Plant Physiol 125, 50–53.

2 Makowka A, Nichelmann L, Schulze D, Spengler K, Wittmann C, Forchhammer K & Gutekunst K (2020) Glycolytic Shunts Replenish the Calvin–Benson–Bassham Cycle as Anaplerotic Reactions in Cyanobacteria. Molecular Plant 13, 471–482.

3 Miao X, Wu Q, Wu G & Zhao N (2003) Sucrose accumulation in salt-stressed cells of agp gene deletion-mutant in cyanobacterium Synechocystis sp. PCC 6803. FEMS Microbiology Letters 218, 71–77.

4 Doello S, Klotz A, Makowka A, Gutekunst K & Forchhammer K (2018) A Specific Glycogen Mobilization Strategy Enables Rapid Awakening of Dormant Cyanobacteria from Chlorosis. Plant Physiology 177, 594–603.

5 Yoo S-H, Lee B-H, Moon Y, Spalding MH & Jane J-L (2014) Glycogen synthase isoforms in Synechocystis sp. PCC6803: identification of different roles to produce glycogen by targeted mutagenesis. PLoS One 9, e91524.

6 Lee K, Doello S, Hagemann M & Forchhammer K (2025) Deciphering the tight metabolite-level regulation of glucose-1-phosphate adenylyltransferase (GlgC) for glycogen synthesis in cyanobacteria. The FEBS Journal 292, 759–775.

7 Ballicora MA, Iglesias AA & Preiss J (2003) ADP-glucose pyrophosphorylase, a regulatory enzyme for bacterial glycogen synthesis. Microbiol Mol Biol Rev 67, 213–225, table of contents.

8 Iglesias AA, Kakefuda G & Preiss J (1991) Regulatory and Structural Properties of the Cyanobacterial ADPglucose Pyrophosphorylases. Plant Physiol 97, 1187–1195.

9 Figueroa CM, Kuhn ML, Hill BL, Iglesias AA & Ballicora MA (2018) Resurrecting the Regulatory Properties of the Ostreococcus tauri ADP-Glucose Pyrophosphorylase Large Subunit. Front Plant Sci 9.

10 Fermont L, Szydlowski N & Colleoni C (2022) Determination of Glucan Chain Length Distribution of Glycogen Using the Fluorophore-Assisted Carbohydrate Electrophoresis (FACE) Method. *Journal of Visualized Experiments (JoVE)*, e63392.

11 Seibold GM, Breitinger KJ, Kempkes R, Both L, Krämer M, Dempf S & Eikmanns BJ (2011) The glgB-encoded glycogen branching enzyme is essential for glycogen accumulation in Corynebacterium glutamicum. Microbiology (Reading) 157, 3243–3251.

12 Wang L, Liu Q, Hu J, Asenso J, Wise MJ, Wu X, Ma C, Chen X, Yang J & Tang D (2019) Structure and Evolution of Glycogen Branching Enzyme N-Termini From Bacteria. Front Microbiol 9.

13 Wang L & Wise MJ (2011) Glycogen with short average chain length enhances bacterial durability. Naturwissenschaften 98, 719.

14 Koch M, Doello S, Gutekunst K & Forchhammer K (2019) PHB is Produced from Glycogen Turn-over during Nitrogen Starvation in Synechocystis sp. PCC 6803. Int J Mol Sci 20, 1942.

15 Wayllace NZ, Valdez HA, Merás A, Ugalde RA, Busi MV & Gomez-Casati DF (2012) An enzyme-coupled continuous spectrophotometric assay for glycogen synthases. Mol Biol Rep 39, 585–591.

16 Eric Holmes & Jack Preiss (1979) Characterization of Escherichia coli B glycogen synthase enzymatic reactions and products. Archives of Biochemistry and Biophysics 196, 436–448.

17 Neumann N, Friz S & Forchhammer K (2022) Glucose-1,6-Bisphosphate, a Key Metabolic Regulator, Is Synthesized by a Distinct Family of α-Phosphohexomutases Widely Distributed in Prokaryotes. mBio 13, e0146922.

18 Kadouche D, Ducatez M, Cenci U, Tirtiaux C, Suzuki E, Nakamura Y, Putaux J-L, Terrasson AD, Diaz-Troya S, Florencio FJ, Arias MC, Striebeck A, Palcic M, Ball SG & Colleoni C (2016) Characterization of Function of the GlgA2 Glycogen/Starch Synthase in Cyanobacterium sp. Clg1 Highlights Convergent Evolution of Glycogen Metabolism into Starch Granule Aggregation1. Plant Physiol 171, 1879–1892.

19 Meléndez R, Meléndez-Hevia E, Mas F, Mach J & Cascante M (1998) Physical Constraints in the Synthesis of Glycogen That Influence Its Structural Homogeneity: A Two-Dimensional Approach. Biophysical Journal 75, 106–114.

20 Plohnke N, Seidel T, Kahmann U, Rögner M, Schneider D & Rexroth S (2015) The Proteome and Lipidome of Synechocystis sp. PCC 6803 Cells Grown under Light-Activated Heterotrophic Conditions *[S]. Molecular & Cellular Proteomics 14, 572–584.

21 Ballicora MA, Iglesias AA & Preiss J (2003) ADP-Glucose Pyrophosphorylase, a Regulatory Enzyme for Bacterial Glycogen Synthesis. Microbiol Mol Biol Rev 67, 213–225.

22 Wegener KM, Singh AK, Jacobs JM, Elvitigala T, Welsh EA, Keren N, Gritsenko MA, Ghosh BK, Camp DG, Smith RD & Pakrasi HB (2010) Global Proteomics Reveal an Atypical Strategy for Carbon/Nitrogen Assimilation by a Cyanobacterium Under Diverse Environmental Perturbations. Mol Cell Proteomics 9, 2678–2689.

23 Kopf M, Klähn S, Scholz I, Matthiessen JKF, Hess WR & Voß B (2014) Comparative Analysis of the Primary Transcriptome of Synechocystis sp. PCC 6803. *DNA Res* **21**, 527–539.

24 Fox J, Kawaguchi K, Greenberg E & Preiss J (1976) Biosynthesis of bacterial glycogen. Purification and properties of the Escherichia coli B ADPglucose:1,4-alpha-D-glucan 4-alpha-glucosyltransferase. Biochemistry 15, 849–857.

25 Kawaguchi K, Fox J, Holmes E, Boyer C & Preiss J (1978) De novo synthesis of Escherichia coli glycogen is due to primer associated with glycogen synthase and activation by branching enzyme. Arch Biochem Biophys 190, 385–397.

26 Cattaneo J, Chambost JP & Creuzet-Sigal N (1978) Combined action of *Escherichia coli* glycogen synthase and branching enzyme in the so-called “unprimed” polyglucoside synthesis. Archives of Biochemistry and Biophysics 190, 85–96.

27 Park J-T, Shim J-H, Tran PL, Hong I-H, Yong H-U, Oktavina EF, Nguyen HD, Kim J-W, Lee TS, Park S-H, Boos W & Park K-H (2011) Role of Maltose Enzymes in Glycogen Synthesis by Escherichia coli▿. J Bacteriol 193, 2517–2526.

28 Boos W & Shuman H (1998) Maltose/maltodextrin system of Escherichia coli: transport, metabolism, and regulation. Microbiol Mol Biol Rev 62, 204–229.

29 Ballicora MA, Sesma JI, Iglesias AA & Preiss J (2002) Characterization of Chimeric ADPglucose Pyrophosphorylases of Escherichia coli and Agrobacterium tumefaciens. Importance of the C-Terminus on the Selectivity for Allosteric Regulators. Biochemistry 41, 9431–9437.

30 Ugalde JE, Parodi AJ & Ugalde RA (2003) De novo synthesis of bacterial glycogen: Agrobacterium tumefaciens glycogen synthase is involved in glucan initiation and elongation. Proc Natl Acad Sci U S A 100, 10659–10663.

31 Hennen-Bierwagen TA, Liu F, Marsh RS, Kim S, Gan Q, Tetlow IJ, Emes MJ, James MG & Myers AM (2008) Starch biosynthetic enzymes from developing maize endosperm associate in multisubunit complexes. Plant Physiol 146, 1892–1908.

32 Tetlow IJ, Beisel KG, Cameron S, Makhmoudova A, Liu F, Bresolin NS, Wait R, Morell MK & Emes MJ (2008) Analysis of protein complexes in wheat amyloplasts reveals functional interactions among starch biosynthetic enzymes. Plant Physiol 146, 1878–1891.

33 Pollock C & Preiss J (1980) The citrate-stimulated starch synthase of starchy maize kernels: Purification and properties. Archives of Biochemistry and Biophysics 204, 578–588.

34 Boehlein SK, Shaw JR, Stewart JD, Sullivan B & Hannah LC (2015) Enhancing the heat stability and kinetic parameters of the maize endosperm ADP-glucose pyrophosphorylase using iterative saturation mutagenesis. Archives of Biochemistry and Biophysics 568, 28–37.

35 Hwang S-K, Hamada S & Okita TW (2006) ATP binding site in the plant ADP-glucose pyrophosphorylase large subunit. FEBS Letters 580, 6741–6748.

36 Baba T, Noro M, Hiroto M & Arai Y (1990) Properties of primer-dependent starch synthesis catalysed by starch synthase from potato tubers. Phytochemistry 29, 719–723.

37 Iglesias AA, Ballicora MA, Sesma JI & Preiss J (2006) Domain Swapping between a Cyanobacterial and a Plant Subunit ADP-Glucose Pyrophosphorylase. Plant Cell Physiol 47, 523–530.

38 Alford JT, Borisova-Mayer M, Mayer C & Forchhammer K (2025) Diverse Metabolic Control of Phosphoglucomutases by Bisphosphorylated Sugars in Heterotrophic Bacteria. Microb Physiol 35, 50–64.

39 Selim KA, Haase F, Hartmann MD, Hagemann M & Forchhammer K (2018) PII-like signaling protein SbtB links cAMP sensing with cyanobacterial inorganic carbon response. Proceedings of the National Academy of Sciences of the United States of America 115, E4861.

40 Wayllace NZ, Valdez HA, Merás A, Ugalde RA, Busi MV & Gomez-Casati DF (2012) An enzyme-coupled continuous spectrophotometric assay for glycogen synthases. Mol Biol Rep 39, 585–591.

41 Vidal R & Venegas-Calerón M (2019) Simple, fast and accurate method for the determination of glycogen in the model unicellular cyanobacterium *Synechocystis* sp. PCC 6803. Journal of Microbiological Methods 164, 105686.

42. Wolstencroft K, Krebs O, Snoep JL, Stanford NJ, Bacall F, Golebiewski M, Kuzyakiv R, Nguyen Q, Owen S, Soiland-Reyes S, Straszewski J, van Niekerk DD, Williams AR, Malmström L, Rinn B, Müller W & Goble C (2017) FAIRDOMHub: a repository and collaboration environment for sharing systems biology research. Nucleic Acids Research 45, D404–D407.

43 Chen X, Schreiber K, Appel J, Makowka A, Fähnrich B, Roettger M, Hajirezaei MR, Sönnichsen FD, Schönheit P, Martin WF & Gutekunst K (2016) The Entner–Doudoroff pathway is an overlooked glycolytic route in cyanobacteria and plants. Proceedings of the National Academy of Sciences 113, 5441–5446.

44 Gründel M, Scheunemann R, Lockau W & Zilliges Y (2012) Impaired glycogen synthesis causes metabolic overflow reactions and affects stress responses in the cyanobacterium Synechocystis sp. PCC 6803. Microbiology 158, 3032–3043.

